# Bottom-up but not Top-down Attention Dominates the Value Representation in the Orbitofrontal Cortex

**DOI:** 10.1101/2021.06.14.448326

**Authors:** Wenyi Zhang, Yang Xie, Tianming Yang

## Abstract

The orbitofrontal cortex (OFC) is essential for value-based learning and decision making. Understanding the attentional modulation of the representation of value in the OFC provides us key information on its functional roles and links the OFC to other cognitive processes. We examined how top-down and bottom-up attention modulates the value encoding in the OFC. Two macaque monkeys were trained to detect a luminance change at a cued location between a pair of visual stimuli, which were over-trained pictures associated with different amount of juice rewards and, thus, different salience. While the monkeys’ behavior and the DLPFC neuronal activities indicated that the monkeys actively directed their attention toward the cued location during the task, the OFC neurons’ value encoding, however, was dominated by the bottom-up attention based on stimulus salience and only reflected the top-down attention weakly. The disassociation between the top-down and bottom-up attention signals in the OFC indicates that the OFC occupies an early stage of value information processing in the brain.

## Introduction

Imagine that you are in a bookstore trying to find an ideal gift for your young child. While you direct your search in the children’s book section, an eye-catching poster of your favorite writer in the bestseller section would divert your attention and lead you to check out the books over there. While task demands or behavior context often lead to internally generated attention, commonly referred as top-down attention, salient stimuli, either based on physical or reward salience, create attention via a bottom-up mechanism (Anderson et al., 2011; Awh et al., 2012; Chelazzi et al., 2013; Connor et al., 2004; Gottlieb et al., 2014; Maunsell, 2004). They may be integrated or compete against each other in the brain and affect decision making.

Previous studies have shown that attention affects value-based decision making by modifying choice options’ subjective value (Krajbich et al., 2010; Towal et al., 2013). Moreover, attention, both in the overt form with gaze shifts and in the covert form without eye movements, has been shown to modulate the representation of value in the orbitofrontal cortex (OFC) (McGinty, 2019; McGinty et al., 2016; Xie et al., 2018), which is a key brain area involved in value-based decision making and adaptive behavior (Ballesta et al., 2020; Hunt et al., 2018; Murray and Rudebeck, 2018; Padoa-Schioppa and Assad, 2006; Wallis and Miller, 2003). In addition, the OFC neuronal activities encoded each choice options’ value alternately, which might be related to attentional shifts (Rich and Wallis, 2016).

However, previous studies on the attentional modulation of OFC’s neuronal activities have not distinguished between the top-down and the bottom-up attention. It is not known how attention from the top-down and the bottom-up sources may interact and modulate the OFC’s value representation. As top-down attention usually reflects the task context better than the bottom-up attention, the former should prevail in well-trained animals’ behavior. One would expect that the attentional modulation of the value representation in the OFC closely resembles that of behavior if the OFC sits in an upstream position in the brain’s executive circuitry. Conversely, a weak top-down but strong bottom-up attentional modulation would indicate that OFC can occupy an early level of value processing. Therefore, comparing how top-down and bottom-up attention modulates the OFC neurons’ responses allows us to gain key insights into the functions of the OFC.

To this end, we trained two macaque monkeys to perform a visual detection task with a Posner cueing paradigm (Posner, 1980). A pair of visual stimuli were presented simultaneously, and a cue indicated which of the stimuli would change its luminance. The monkeys had to direct their attention toward the cued stimulus to detect the luminance change. In addition, the stimuli were well trained shapes that were associated with different amounts of reward. They acquired different levels of salience through over-training and generated attention likely via a bottom-up mechanism. Both the monkeys’ behavior and the lateral prefrontal cortex (LPFC) neuronal activities indicated that the monkeys directed their attention according to the cue while the stimulus salience only had a minor influence on the behavior. In contrast, the OFC’s value encoding was only weakly modulated by the top-down attention. The value of the more salient stimulus was predominantly represented in the OFC, regardless of whether it was the target of the top-down attention. The results indicate that the OFC maintains a stable value representation independent from behavior contexts and suggest that the OFC operates at an early stage of value information processing in the brain.

## Results

### Behavioral Task and Subject Performance

We trained two monkeys to perform a visual detection task (**Figure 1a**). In this task, the monkeys were asked to detect when one of the two stimuli changed its luminance after a random delay. The stimulus with the luminance change (target) was cued by a frame that was presented on the side opposite to the change location 200 ms before the onset of the stimulus pair. The cue was valid in 90% (monkey D) or 80% (monkey G) of the trials. In the remaining trials (invalid-cue trials), the target stimulus was on the same side of the frame. The monkeys reported the luminance change by making a saccade toward an eye movement target above the fixation spot.

**Figure 1.**
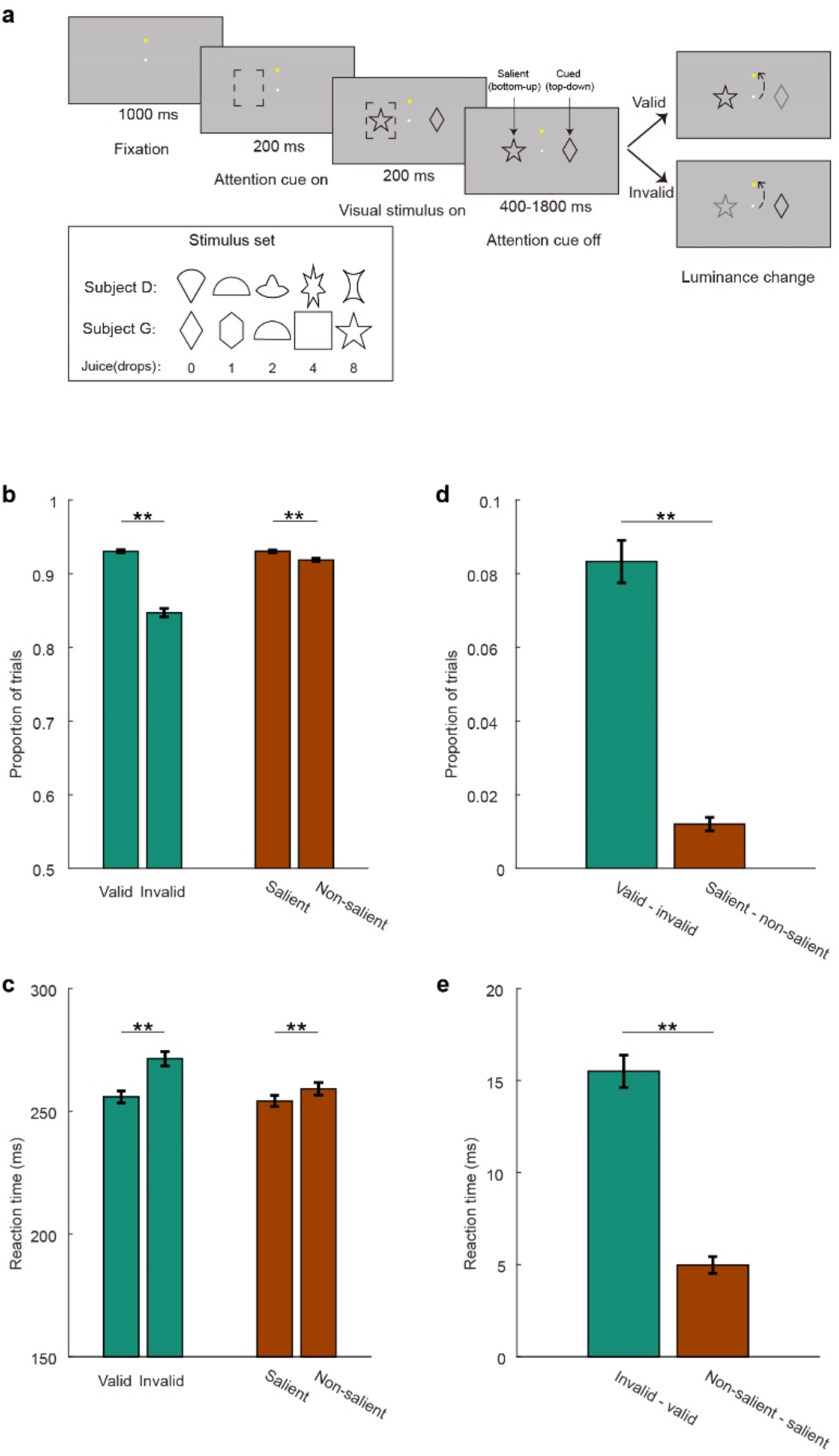
Behavioral Task and Performance. **a.** The monkeys had to detect when one of two stimuli changed its luminance. The location of the luminance change was indicated by a cue (monkey D: 90% valid; monkey G: 80% valid) that appeared on the side opposite to the luminance change. Invalidly cued trials had the luminance change on the same side of the cue. Subjects reported the luminance change by making an eye movement to the eye-movement target located above the fixation point. The stimuli were associated with different amounts of juice reward (inset). Random pairings of stimuli were selected for each trial. The stimulus with larger associated reward is referred as the salient stimulus (star in the example), and the stimulus where attention should be direct to is the cued stimulus (diamond). The computer randomly selected one stimulus from the pair and delivered its associated reward for a correct response. **b.** The proportion of correct responses when the cue was valid and invalid, and when the luminance change occurred at the salient and the non-salient stimulus. **c.** The reaction time of detected luminance changes when the cue was valid and invalid, and when the luminance change occurred at the salient and the non-salient stimulus. **d.** The improvement of performance due to cue validity (green) and the salience (brown). **e.** The improvement of reaction time due to cue validity (green) and the salience (brown). The error bars are SEM across sessions (monkey D: n=71; monkey G: n=100), ** denotes p<0.001.

The stimulus set includes 5 pictures for each animal. Each picture was associated with a juice reward of different size (**Figure 1a**, inset). We initially trained the monkeys with single-stimulus trials, in which the frame appeared on the side opposite to the single cue. Its associated reward would be delivered for a correct response. On the double-stimulus trials, the reward delivered after a correct response was randomly chosen between the rewards associated with the two stimuli. Therefore, in the single-stimulus trials the monkeys were certain of the reward they could get from a correct response, while in the double-stimulus condition the reward outcomes were only 50% certain.

For the ease of the discussion, between the two stimuli presented in each trial, we refer the stimulus that is cued to have a luminance change as the cued stimulus, its value as the cued value (CV). The other stimulus is referred as the uncued stimulus, and its value as the uncued value (UCV). As the monkeys were over-trained with this task and were familiar with the stimuli, the stimuli acquired different levels of salience due to their reward associations. Between the two stimuli presented in each trial, we refer the stimulus associated with a larger reward as the salient stimulus, and the other one the non-salient stimulus. Their values are termed the salient value (SV) and the non-salient value (NSV). The SV and the NSV are the same in trials with two of the same stimuli.

After the training, the monkeys performed the task well. When the cue was valid, they were more accurate (valid: 93.0%, SEM=0.2%; invalid: 84.7%, SEM=0.6%. p<<0.001, two-tailed Wilcoxon signed rank test) and reacted faster (valid: 256 ms, SEM=2 ms; invalid: 271 ms, SEM=3 ms. p<<0.001, two-tailed Wilcoxon signed rank test) (**Figure 1b, c**, green bars). The monkeys understood the meaning of the cues and took advantage of them by directing their attention appropriately.

The stimulus salience also improved the monkeys’ performance, although to a much lesser degree. When the more salient stimulus changed luminance, the monkeys were more accurate (salient: 93.1%, SEM=0.2%; non-salient: 91.8%, SEM=0.2%. p<<0.001, two-tailed Wilcoxon signed rank test) and reacted faster (salient: 254 ms, SEM=2 ms; non-salient: 259 ms, SEM=3 ms. p<<0.001, two-tailed Wilcoxon signed rank test) (**Figure 1b, c**, brown bars). These behavior improvements caused by the salience were however much smaller than those caused by the cue (two-sided Wilcoxon signed rank test, p<<0.001 for both accuracy and reaction time, **Figure 1d, e**). On average, paying attention to the salient stimulus does not bring any behavior benefits to the monkey or provide information about the expected reward. Correspondingly, the top-down attention generated by the cue clearly dominated the monkeys’ behavior.

### LPFC Population Encodes Spatial Attention Shifts

The behavior analyses revealed that the animals directed their spatial attention mostly according to the cue during this task. We can also use neuronal activities to determine the animals’ spatial attention location, which can provide us more detailed information on how the animals directed their attention during the task. The activity of the lateral prefrontal cortex (LPFC) neurons can be used to decode the location of top-down spatial attention (Tremblay et al., 2015a, 2015b). Therefore, we recorded the activity of 422 LPFC single units (monkey D: 255; monkey G: 167) from Walker’s areas 8a, 8b, 46d and 46v (**Supplementary Figure 1**) and used those signals to decode the location of attention.

One important feature of our task is that the frame used as the cue was displayed on the side opposite to where the attention needed to be directed. The onset of the frame captured the monkeys’ attention via a bottom-up process. Afterwards, the monkeys had to shift the attention to the opposite side where the luminance change was expected. This attention shift was captured by many LPFC neurons that encoded spatial attention location. In **Figure 2a** we show an example of one such neuron. The neuron had larger responses when the frame appeared to the right of the fixation point (blue trace), but only transiently. Once the stimuli appeared its response was much greater in trials that had the cue frame on the left and the cued stimulus on the right (red trace). The neuron’s responses reflected the attention shift from the side of the frame to the opposite side where the luminance change was expected.

**Figure 2.**
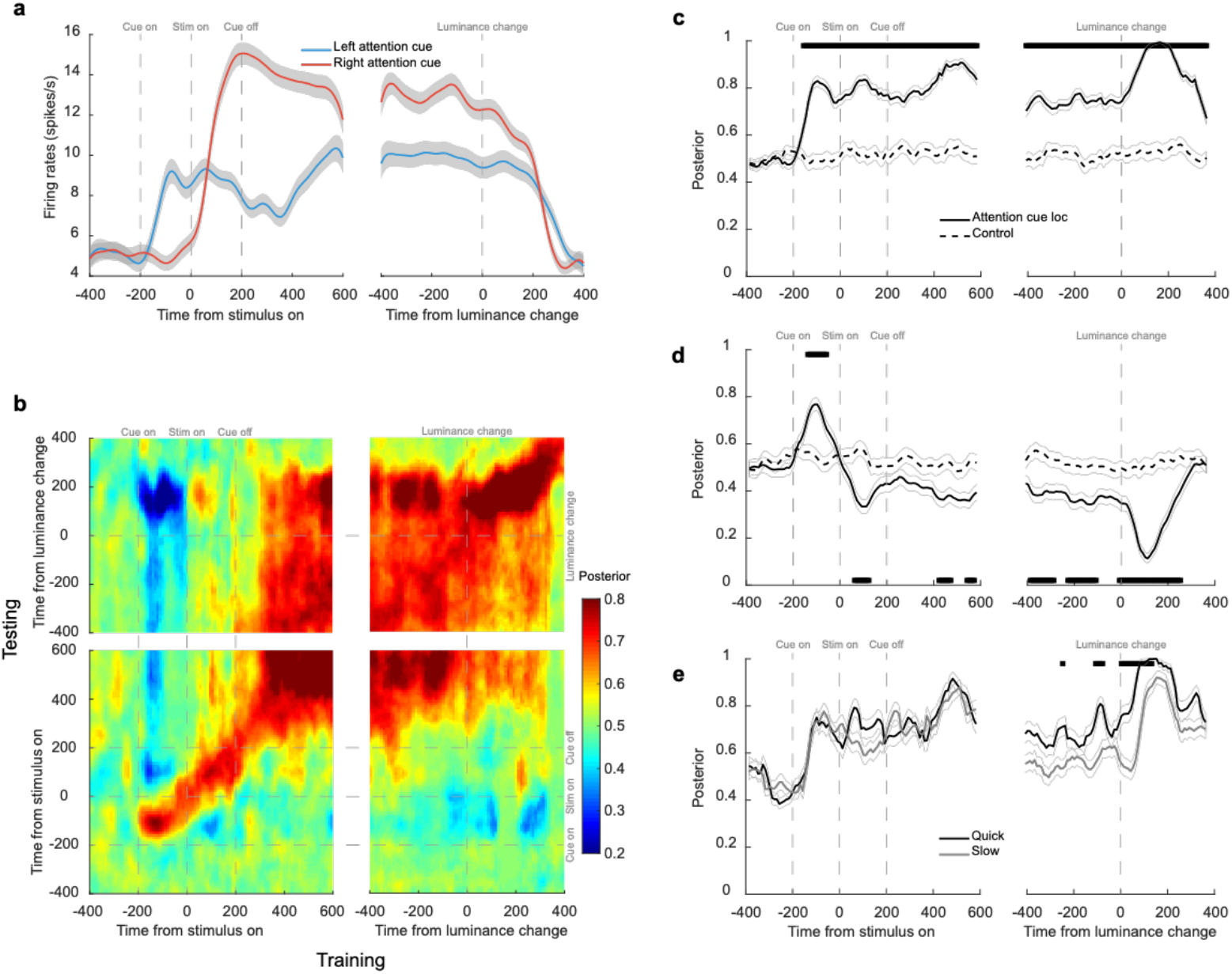
LPFC encoded top-down spatial attention. **a.** The responses of an example LPFC neuron in double-stimulus trials. Red line indicates trials when the cue was on the left side and blue line indicates trials when the cue was on the right side (blue line). The shaded areas around each line denote SEM across trials. **b.** The posterior probability of the location of the top-down spatial attention across trials decoded from LPFC pseudo neuronal population ensemble activities. Both the training and the testing of the LDA decoder used sliding windows of 25 ms at 10 ms steps. **c.** The posterior probability of the location of the top-down attention from the decoder that was trained and tested with the LPFC neuronal responses at the same time (the diagonal line in **b**). Significance was assessed with two-tailed paired t-tests (p<0.01 with FDR corrections for multiple comparisons), testing against the data with shuffled cue location labels. The black segments at the top indicate when the actual population performed significantly better than the control. Thin grey lines represent SEM across trials. **d.** Same as **b** except that the decoder was trained with the mean activities at 50 to 200 ms before the stimulus onset. The black segments at the bottom indicate when the performance was significantly lower than the control (p<0.01 with FDR corrections for multiple comparisons). **e.** The posterior probability of the location of the spatial attention in the fast (black line) and the slow trials (grey line). The trials were split according to the median reaction time in each session. Significance was assessed with two-tailed paired t-tests (fast versus slow, at p<0.01 with FDR corrections for multiple comparison).

Population analyses confirmed that the LPFC neurons encoded the spatial attention location and reflected the attentional shift. To pool neurons with different spatial selectivity together, we considered the whole population of recorded neurons as a high-dimensional representation of task related variables (Mante et al., 2013; Rich and Wallis, 2016; Rigotti et al., 2013) and trained a linear discriminant analysis (LDA) algorithm with the pseudo neuronal ensemble to decode the frame location. The pseudo neuronal ensemble was composed in such a way that each neuron had the same number of left and right frame trials (see Methods). The decoder based on the LDA provided the posterior probabilities of the frame location throughout the task.

We trained and tested the decoders with LPFC neurons’ responses using a sliding window (size: 25 ms, step: 10 ms) and plotted the decoders’ performance as a heat map. When the decoder was trained and tested with the responses at the same time point, indicated by the diagonal in **Figure 2b**, the cue location can be reliably decoded based on the LPFC population responses shortly after the onset of the attention cue (**Figure 2c**, left) and until after the luminance change and the animal’s response (**Figure 2c**, right). During the period from ~200 ms after the stimulus onset until after the luminance change, the encoding of the attention location was stable so that the cross-temporal decoding performance was good (**Figure 2b**). Furthermore, the cross-temporal decoder’s performance revealed the attention shift after the shape onset. We trained the decoder with the neuronal responses 50 to 200 ms before the stimuli onset when the frame was the only stimulus on the screen, and tested its performance using the responses at the other time points in a trial (**Figure 2d**). The decoder initially performed well above the shuffled level during a short period after the frame onset, but its performance quickly dropped to below the shuffled level after the stimulus onset and reached to its minimum after luminance change. Lower-than-shuffled performance indicates that the attention location shifted to the side opposite to where it was when the decoder was trained. This attention shift also appears in **Figure 2b** in the blue regions – indication of the performance below the chance level –where the decoders were trained and tested at time points when the frame was presented and when the stimuli were presented.

Finally, the encoding of the attention location in the LPFC correlated with the behavior performance. We divided the trials by the monkeys’ reaction time into two halves and tested the decoder’s performance in each half of trials. The decoder performed significantly better before the onset of the luminance in trials when the monkeys reacted faster to the luminance change, suggesting that the attention was more likely to be allocated correctly in those trials (**Figure 2e**).

### OFC Responses were Dominated by the Salient Value

After we verified that the monkeys directed their attention to the proper location during the task, we investigate how the OFC neurons’ value encoding might be modulated by the attention.

In the current study, there were two sources of attention. One was induced by the frame cue, which indicated the location of the luminance change. The attention was first directed at the location of the cue itself, presumably due to the onset of the cue and via a bottom-up process, but the monkeys then directed their attention to the opposite side for detecting the luminance change via a top-down process. The other source of the attention was caused by the stimuli themselves. The stimuli were associated with different amounts of juice or water reward. After months of training, they acquired different levels of salience because of their reward association.

We recorded the activities of 357 OFC neurons (monkey D: 220; monkey G: 137) from Walker’s areas 11 and 13 (Supplementary Figure 1). Many of them encoded stimulus value. An example neuron is shown in **Figure 3a,d,g,j**. We first used trials in which two stimuli with the same value to measure the neurons’ value selectivity. In these trials, the monkeys were certain of the reward they would get for a correct response, and our previous study indicated that the OFC neurons responded similarly to a pair of the identical stimulus and the same stimulus presented alone (Xie et al., 2018). The example neuron’s responses reflected the value of the stimuli; its responses were larger when the value was larger (**Figure 3a**). To study the neuron’s responses to a pair of distinct stimuli, we first grouped the trials by the SV and observed a very similar pattern (**Figure 3d**). The neuron’s responses reflected the SV stably during the stimulus period well until after the luminance change. We calculated the neuron’s firing rates between the stimulus onset and the luminance change and plotted them against the SV (**Figure 3g**). The responses to the SV were similar to the responses when a pair of the same stimuli was presented. The non-salient stimulus was ignored. In comparison, when we looked at how the attention cue affect the value encoding by grouping the trials by the CV and the UCV, the neuron exhibited similar responses, indicating both the CV and the UCV contributed to the neuron’s responses similarly (**Figure 3j**). The results suggested that the salience but not the cue dominated the example neuron’s responses.

**Figure 3.**
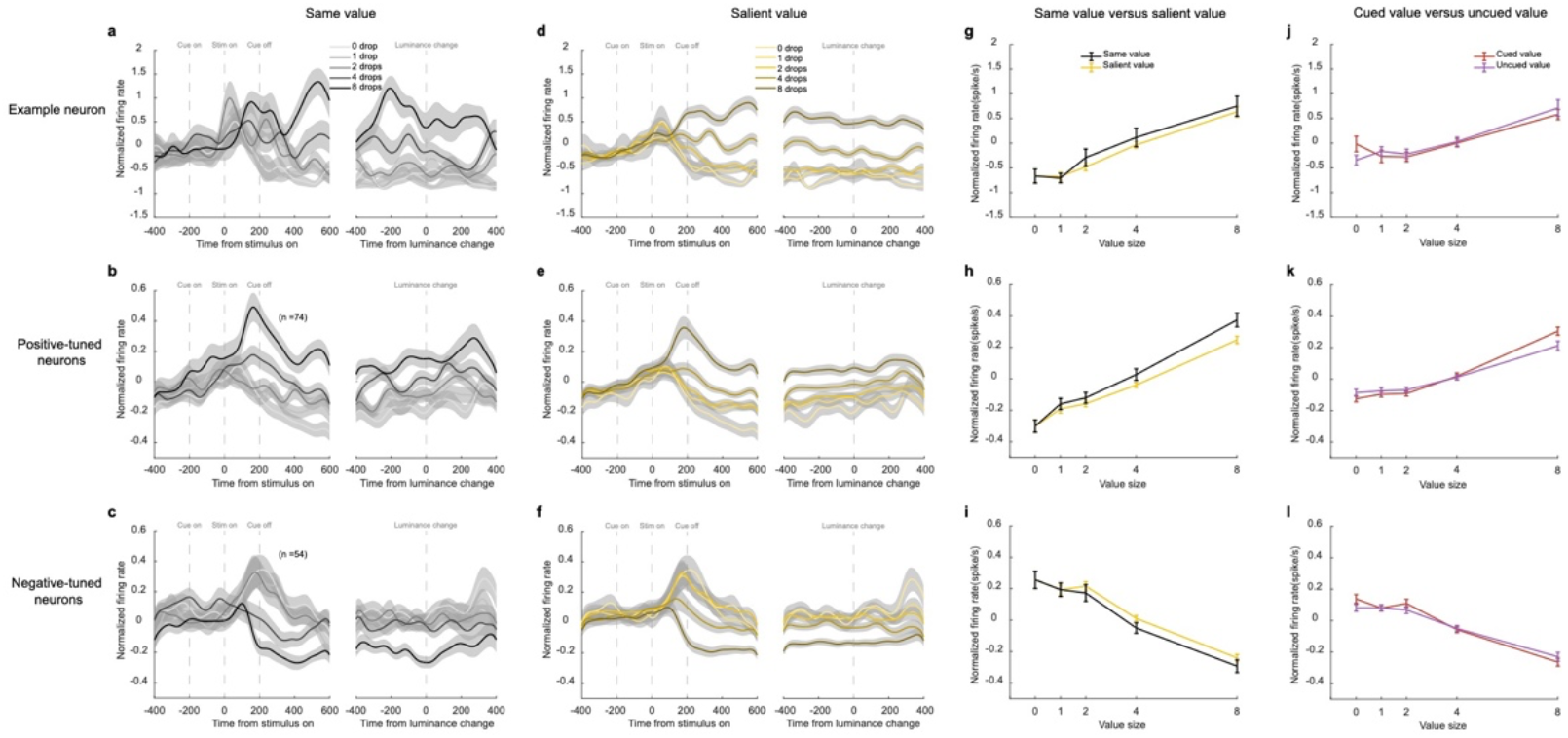
Attentional modulation of the OFC responses. **a.** The responses of an example OFC neuron in trials when the same rewards were the associated with the two stimuli, aligned to the stimulus onset (left) and the luminance change (right). The reward sizes are indicated by different shades of grey. Shading areas indicate SEM across trials. **b.** The population response of the positively tuned OFC neurons (n=74) in trials when the same rewards were the associated with the two stimuli. Shading areas indicate SEM across neurons. **c.** Same as **b**, but for the negatively tuned OFC neurons (n=54). **d.** The responses of the example neuron with the trials grouped by the SV. The reward sizes are indicated by different shades of yellow. Shading areas indicate SEM across trials. **e.** The responses of the positively tuned OFC neurons with the trials grouped by the SV. The reward sizes are indicated by different shades of yellow. Shading areas indicate SEM across neurons. **f.** Same as **d**, but for the negatively tuned OFC neurons. **g.** The average responses during the period between the stimulus onset and the luminance change. The black trace is based on the trials with the same-reward stimulus pairs. The yellow trace is based on the trials grouped by the SV. The error bars indicate SEM across trials. Two-way ANOVA (group: F_1,413_=0.8, p=0.3729; value: F_4,413_=40.26, p<<0.001). **h.** The average responses of the positively tuned OFC neurons during the period between the stimulus onset and the luminance change. The black trace is based on the trials with the same-reward stimulus pairs. The yellow trace is based on the trials grouped by the SV. The error bars indicate SEM across neurons. Two-way ANOVA (group: F_1,730_=6.57, p=0.0105; value: F_4,730_=103.35, p<<0.001). **i.** Same as **h**, but for the negatively tuned OFC neurons. Two-way ANOVA (group: F_1,530_=1.52, p=0.2189; value: F_4,530_=58.44, p<<0.001). **j.** Same as **g**, but trials are grouped by the CV (red) and the NCV (purple). Two-way ANOVA (group: F_1,690_=0, p=0.9889; value: F_4,690_=21.04, p<<0.001). **k.** Same as **h**, but trials are grouped by the CV (red) and the NCV (purple). Two-way ANOVA (group: F_1,730_=0.07, p=0.7904; value: F_4,730_=105.8, p<<0.001). **l.** Same as **i**, but trials are grouped by the CV (red) and the NCV (purple). Two-way ANOVA (group: F_1,530_=0.6, p=0.4385; value: F_4,530_=86.16, p<<0.001)

The OFC population exhibited the same trend. Among the 357 OFC neurons from which we recorded, 128 neurons were selective to value (**Supplementary Table 1**). We further divided these neurons into positively and negatively tuned based on their value tuning. We used trials with a pair of the same-value stimuli to remove value uncertainty when measuring the neurons’ value selectivity. The positively tuned neurons had larger responses when the values were higher (**Figure 3b**, 74 out of 128 value-tuned neurons) and the negatively tuned ones had the opposite tuning (**Figure 3c**, 54 out of 128 value-tuned neurons). Both groups of neurons encoded the SV stably during the stimulus period (**Figure 3e,f**). Their firing rates sorted by the SV were similar to when a pair of the stimuli with the same value as the SV were presented (**Figure 3h,i**), and their responses did not distinguish between the CV and the NCV (**Figure 3k,l**).

We further quantified the contribution of the SV and the NSV to the OFC neuronal responses with a linear regression model that contained both value variables as well as a binary term that indicated the frame location. We calculated the coefficient of partial determination (CPD) to measure how much variance was explained by each variable (see Methods). The average CPD of the SV rose significantly above the baseline shortly after the stimulus onset (**Figure 4a**). In contrast, the CPD of the NSV remained low and not significantly different from the baseline in most time bins. The paired t-tests that compared the CPDs between the SV and the NSV indicated that the OFC neurons encoded the SV much better in almost all the time bins after the stimulus onset (p < 0.005 with FDR correction for multiple comparisons). The frame location was transiently encoded when the frame was presented and after the luminance change but was not significantly encoded during most of the stimulus presentation period.

**Figure 4.**
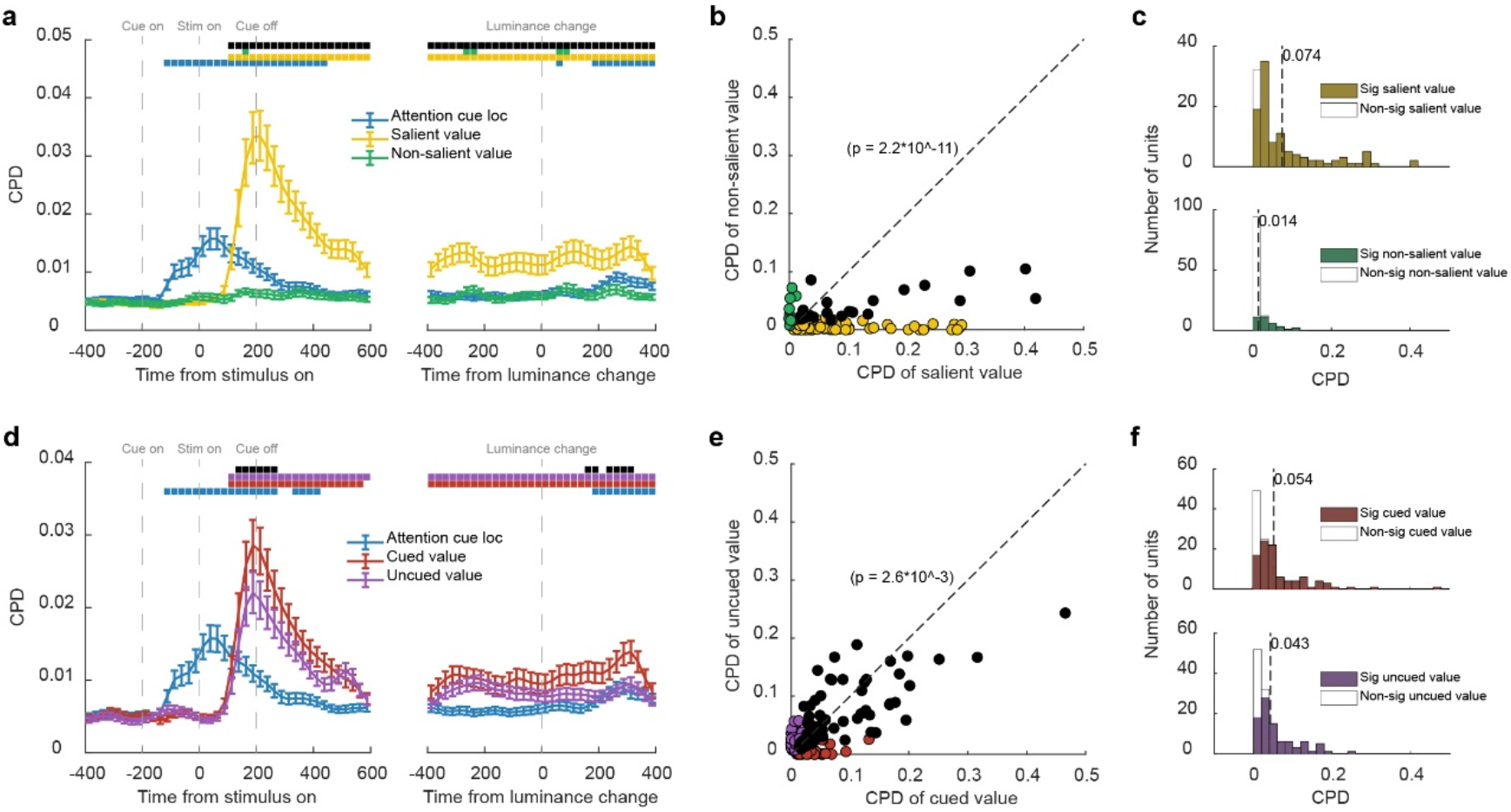
OFC neuronal responses dominantly encoded the salient value. **a.** Time course of average coefficients of partial determination (CPD) for the SV and the NSV. Significance was assessed with two-tailed paired t-tests (p < 0.005, with FDR corrections for multiple comparison) comparing to a baseline computed with the average CPD between 0 and 200 ms before the cue onset and across different regressors. The blue, yellow and green bars at the top indicate the significant CPDs of the cue location, the SV, and the NSV, respectively, and the black bar indicates the significant difference between the SV and the NSV. The error bars indicate SEM across neurons. **b.** The CPDs of the SV against the CPDs of the NSV for individual neurons. The yellow data points are the neurons with non-zero CPDs for the SV only, the green data points are the neurons with non-zero CPDs for NSV only, and the black data points are the neurons that encoded both the SV and the NSV (p<0.05). Paired t-test was conducted to compare between the mean CPDs of the SV and the NSV. **c.** Top: the distribution of the CPDs for the SV of the value-selective neurons. Bottom: the distribution of the neurons’ CPDs for the NSV. Vertical dashed lines indicate the mean. Filled bars indicate significant neurons. **d.** Same as **a**, but for the CV and the UCV. **e.** Same as **b**, but for the CV and UCV. **f.** Top: the distribution of the CPDs for the CV of the value-selective neurons. Bottom: the distribution of the neurons’ CPDs for the NCV. Vertical dashed lines indicate the means. Filled bars indicate significant neurons.

This trend is also observed at the level of individual neurons. We calculated each neuron’s average responses during the stimulus period and used them to determine the CPD for the SV and the NSV for each value-selective neuron (**Figure 4b**). The data points of the neurons spread along the axis of the CPD for the SV (SV: 105/357 significant, p<0.05, linear regression). Their CPDs for the NSV, on the other hand, sat within a small range (NSV: 34/357 significant, p<0.05, linear regression). The distribution of the neurons’ CPD for the SV were much more extended in the range than that for the NSV, and the mean of the CPDs for the SV was larger than that of the UCV (p=2.2×10^-11, paired t-test) (**Figure 4c**).

By contrast, when we regressed the OFC neuronal responses against the CV and the UCV, we obtained overall similar CPDs, suggesting that the CV and the UCV contributed similarly to the responses (**Figure 4d**). Other than a short period after the stimulus onset and another one after the luminance change, the difference between the CV and the UCV’s CPDs were not significant during most of the stimulus period. During the period when the CPD of the CV was larger, there were extra visual inputs, the frame initially and the luminance change in the end, which might serve as a bottom-up attention signal. When we plotted each neurons’ CPDs for the CV and against those for the UCV, they tended to spread more closely to the diagonal than their CPDs for the SV plotted against the NSV (**Figure 4e**). The CPDs for the CV were slightly but significantly higher than those for the NCV (p=0.003). These results indicate that the cue and the top-down attention only weakly and transiently modulated the neurons’ value tuning.

Finally, to confirm that the weak top-down attentional modulation was not due to a washout effect by the many trials in which the CV and the UCV had similar values, we studied the trial conditions in which we would expect the largest modulation on the neurons’ responses by the top-down attention. According to our previous study (Xie et al., 2018), the cue-induced modulation might be most evident when the cue directed the attention toward the non-salient picture when the SV was the greatest (8 drops of juice). In this condition, the cue shifted the attention away from the most salient picture, which captured the monkeys’ attention by default. This would provide us an opportunity to observe any potential modulation of the neurons’ responses by the cue in addition to the pictures’ salience. Therefore, we studied all trials in which one of the two stimuli was associated with 8 drops of juice. We compared the neurons responses when the cue directed the attention toward the 8-drops-of-juice cue (CV=8) and when the cue directed attention away from it (CV=0, 1, 2, 4, or 8). Again, we looked at the positively- and negatively tuned OFC neurons separately. When the attention was on the 8-drops-of-juice cue, the OFC neurons ignored the value of the less salient stimulus (linear regression, H_0_: slope=0: p=0.556, positively tuned neurons; p=0.4455, negatively tuned neurons). The responses were highest for the positively tuned neurons and lowest for the negatively tuned ones, both similar to their responses when only 8-drops-of-juice were presented (**Figure 5a,b**). When the top-down attention was directed away from the 8-drops-of-juice cue, it lowered the responses of positively tuned neurons slightly (F_720,1_=6.68, p=0.01) but failed completely to increase the responses of the negatively tuned neurons (F_530,1_=0.96, p=0.3271).A complete shift of responses toward the less salient stimulus would have produced responses similar to those when the less salient stimulus was presented alone (black trace in **Figure 5a,b**). The neuronal responses were still similar to the responses to the more salient stimuli, the 8-drops-of-juice stimulus. Attending to the less salient stimulus did not abolish the dominant representation of the SV by the OFC.

**Figure 5.**
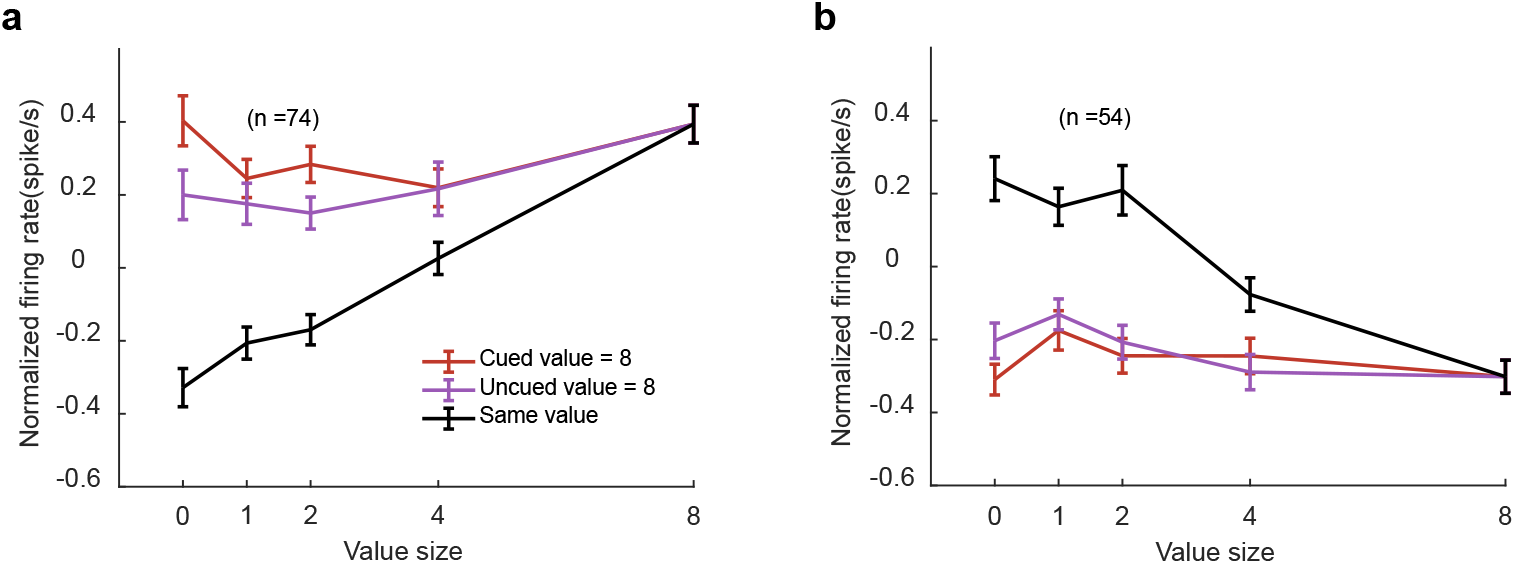
Top-down attention failed to switch the value encoding in the OFC to the non-salient stimulus. **a.** The positively tuned OFC neurons’ responses to an 8-drops-of-juice stimulus (SV) paired with stimulus associated 0,1,2,4, or 8 drops of juice (NSV). The red line denotes trials when the top-down attention was toward the 8-drops-of-juice stimulus, and the purple line denotes trials when the top-down attention was away from the 8-drops-of-juice stimulus. A two-way ANOVA (attention location: F_720,1_=6.68, p=0.01; value: F_720,4_=3.95, p=0.0035) indicates a subtle but significant top-down attentional modulation. A complete switch of the value encoding to the NSV would produce responses close to the black line, which are the neurons’ responses to the trials of a pair of stimuli with the same reward. All error bars indicate SEM across neurons. **b.** Same as **a**, but for the negatively tuned OFC neurons. Two-way ANOVA with group (CV=8/UCV=8) and value was performed (group: F_530,1_=0.96, p=0.3271; value: F_530,4_=2.85, p=0.0235).

## Discussion

Here, we demonstrated that while both the behavior and the DLPFC neuronal responses indicated that the monkeys directed their attention to the cued location, the OFC neuronal responses were nevertheless dominated by stimulus salience and only weakly affected by the top-down attention. As top-down attention often accompanies the form of decision, many brain areas implicated in decision making possess a strong top-down attention signal (Bisley and Goldberg, 2003; Fecteau et al., 2004; Tremblay et al., 2015a). The lack of strong top-down attention signal in the OFC indicates that the OFC occupies an early stage of value information processing in the brain.

Our behavior paradigm allowed us to tease apart bottom-up attention, top-down attention, reward, decision making, and eye movements. First, the frame that served as the top-down attention cue was opposite to the location that needed to be attended to. The monkeys switched their attention from the frame to the cued location, and any attentional effects we observed at the cued location before the luminance change was not due to the bottom-up attention. Second, the monkeys had to report their decision by making a saccade to a target above the fixation point. The motor preparation signals were thereby dissociated from the attention signals. Lastly, the reward that the monkey received were randomly chosen between the two stimuli. Neither the cued value nor the salient value was the expected reward outcome. Moreover, no decisions were guided by the values of the stimuli, so potential attention shifts accompanying the value-based decision-making process was minimized.

Here, we most used the value saliency as a proxy for studying bottom-up attention. The bottom-up attention considered in this study is an associated salience acquired through extensive training and may be distinct from bottom-up attention based on physical salience. It may involve a subcortical circuitry that includes superior colliculus, pulvinar, ventral midbrain, and amygdala (McFadyen et al., 2019; Takakuwa et al., 2017). However, physical salience also produced bottom-up attention signals that we observed in the OFC. For example, the onset of the frame, although by itself, was valueless, provided a strong bottom-up signal. It was reflected in the OFC neuronal activity (**Figure 4a, d**, blue trace), but only briefly, reflecting the transient nature of this bottom-up signal. Furthermore, the luminance change, although subtle, might also induce the bottom-up attention toward the cued stimulus. This luminance change was similar to the visual perturbation used in our previous study, which led to a modulation of the OFC’s value encoding (Xie et al., 2018). We also observed a slightly stronger representation of the cued stimulus than that of the uncued after the luminance change (**Figure 4d**). It could be explained as a bottom-up attention effect. This modulation effect could not be explained by any motor plans, as the planned eye movements were in a direction orthogonal to the attended and the unattended stimuli.

The dominance of the bottom-up modulation, which was almost all-or-none, is in stark contrast to how neurons in the visual cortex are modulated by attention (Maunsell, 2015). The responses of the visual neurons to multiple stimuli presented in their receptive field can often be well described with a normalization model that performs a weighted average of the responses to each stimulus presented alone. Attention modulates how the responses are weighted by adding bias toward the attended stimulus. In addition, the neuron’s stimulus preference plays an important role. Whether attended or not, a preferred stimulus presented in a neuron’s receptive field has a significant influence on its responses (Ni et al., 2012). The OFC neurons, however, only encode the value of the most salient stimulus, even for the negatively tuned neurons, which prefer the non-salient stimulus. The less salient stimuli are ignored by the OFC neurons, and top-down attention cannot help very much.

Our results imply that the OFC does not integrate information from multiple simultaneously presented stimuli, as we showed that the less salient stimuli are ignored by the OFC. Although the representation of the salient value may be interpreted as a decision signal in the OFC, it has been demonstrated that the OFC neurons do not accumulate information from sequentially presented stimuli in a value-based decision-making task (Lin et al., 2020). The lack of information integration across space or time suggests that the OFC’s role in value processing is more analogous to that of the early visual cortices in visual processing. Accordant with this view, a recent study showed that micro-stimulation in the OFC affected value-based decision making (Ballesta et al., 2020).

For now, we can only speculate the functional significance of the fact that the OFC maintains a representation of the salience signal independent of the top-down attention. Such a representation may be necessary for the brain to evaluate the potential targets toward which the top-down attention could be directed. The OFC may provide the basis for this computation. Future experiments could be designed to investigate this hypothesis.

## Funding

This work was supported by the National Natural Science Foundation of China (Grant No. 31771179), Shanghai Municipal Science and Technology Major Project (Grant No. 2018SHZDZX05), and the Strategic Priority Research Program of Chinese Academy of Science (Grant No. XDB32070100).

## Acknowledgements

We thank John Maunsell for his comments during the preparation of the manuscript. We also thank Ruixin Su, Wei Kong, Lu Yu, and Tian Qiu for their help in all phases of the study. The authors declare no competing financial or nonfinancial interests.

## Methods

### Subjects

Two male rhesus monkeys (*Macaca mulatta*) were used. They weighed 9.3 kg (subject D) and 6.8 kg (subject G) at the beginning of the training. All experimental procedures were approved by the Animal Care Committee of Shanghai Institutes for Biological Sciences, Chinese Academy of Sciences (Shanghai, China).

### Behavioral task and materials

Head restrained monkeys were seated in a primate chair facing a 23.6-inch computer monitor at 60 cm away from their eyes. The center of the screen was adjusted to align to the midpoint of the two eyes. Behavioral tasks were run with the MATLAB based software MonkeyLogic (Asaad et al., 2013). Subjects’ eye position and pupil dilation were tracked with an infrared oculometer system at a sampling rate of 500Hz (EyeLink 1000). Juice was delivered by a computer-controlled solenoid.

We trained two monkeys to perform a visual detection task (**Figure 1a**). The subjects had to hold their gaze at a central fixation point on the screen within a 3° wide window. After the monkeys maintained fixation for 0.5 s, a yellow saccade target was presented 6.0° (monkey D) or 7.6° (monkey G) above the fixation point. The saccade target remained on the screen until the end of the trial. After 0.5 s, a square frame (attention cue) 2.4° wide was presented on the left or right (subject D: 7.0°; subject G: 7.6°) of the fixation point. After another 0.2 s, either one or two visual stimuli of 2.4° size appeared on the left, the right, or both sides (monkey D: 7.0°; monkey G: 7.6 °) of the fixation point. The stimulus locations were rotated slightly (23° counter-clockwise) for monkey G to minimize its left-right bias. The attention cue disappeared 0.2 s after the stimulus onset. The animals were required to maintain their fixation until they detected the target stimulus changed its luminance, upon which they needed to saccade to the target between 100 to 400 ms after the luminance change to receive a reward. The onset latency of the luminance change was randomly chosen from an exponential distribution (tau = −2.5 s, cut off at 1.4 s) plus 0.4 s.

In trials with one stimulus, the attention cue indicated that the upcoming stimulus would appear on the opposite side of the fixation point (always valid). In trials with two visual stimuli, the attention cue indicated the stimulus on the opposite side of the fixation point was the target stimulus. The attention cue was valid in 90% (monkey D) or 80% (monkey G) of trials. In the remaining trials, the stimulus on the same side of the fixation point was the target stimulus. Monkey D was trained to detect the luminance change of both the valid and the invalid cue. The monkey G was trained to maintain the fixation when the luminance change occurred at the invalid cue and only respond to the change at the valid cue.

The visual stimuli were associated with different amounts of juice that monkey might get for a correct response. There were five stimuli for each monkey, each was associated with 1 small drop (0.033 ml), 1, 2, 4, and 8 standard drops (0.10, 0.16, 0.29, and 0.55 ml) of juice, respectively. For convenience, we label the stimulus with a small drop of juice as 0 standard drops. During the initial training, only one stimulus was presented on the screen, and the monkeys were rewarded with its associated juice amount if they detected the luminance change correctly. Afterwards, we introduced double-stimulus trials. When two stimuli were presented, one of the stimuli was randomly selected and monkey would receive the reward associated with that stimulus.

The single-stimulus and double-stimulus trials were interleaved in blocks. The stimuli in each trial were randomly selected from the stimulus set, and their locations (left or right) were counterbalanced.

### Surgery and MRI

Before the behavioral training, both monkeys received a chronic implant of a titanium headpost. After two-month recovery, they were trained to perform the behavioral task until they achieved satisfactory performance. Then, the monkeys received structural Magnetic Resonance Imaging (MRI) scans for us to determine the recording chamber location. After the chamber implant surgery, a manganese-enhanced MRI scan was conducted to verify the chamber placement. The scans were carried out in a Siemens 3T scanner.

During the surgery, the monkeys were sedated with ketamine hydrochloride (10 mg/kg), and anesthesia was then induced and maintained with isoflurane gas (1.5-2%, to effect). Body temperature, heart rate, blood-oxygen concentration, and expired CO_2_ were monitored throughout the surgical procedures.

### Neuronal recordings

A 2 cm * 1.5 cm chamber was implanted on the surface of left (subject G) or right (subject D) prefrontal cortex, centered 31.5 (subject G) or 27.5 (subject D) mm anterior along the anterior-posterior coordinate plane (Supplementary Figure 1). We recorded extracellular single-unit activities with tungsten microelectrodes (FHC: 0.3-2 MΩ; AlphaOmega: 0.5-3 MΩ). Each electrode was driven by an independent microdrive (AlphaOmega EPS) through a stainless-steel guide tube. The guide tube was placed within a grid with holes 1 mm apart. The depth of the penetration was confirmed by the transitions between gray and white matter. At most four electrodes were lowered at a time. Neuronal signals were recorded with an AlphaOmega SnR system at a sampling rate of 44 kHz. We recorded neurons from the OFC and the LPFC. The OFC recording locations were between the lateral and medial orbital sulci in Walker’s areas 11 and 13. The LPFC neurons were recorded from the area rostral to the arcuate sulcus (8a and 8b), including both the dorsal and the ventral bank of the principal sulcus (46d and 46v).

Voltage signals of putative single neurons were isolated offline manually with Plexon Offline Sorter (Plexon, Dallas, TX). Neurons with poor isolation or with lower than 1 Hz response rate were excluded. There were no additional selection criteria for neurons.

### Behavioral analyses

All the behavior analyses were based on the monkeys’ performance during the recording sessions.

### Accuracy and reaction time

To calculate behavior accuracy in **Figure 1b**, we measured hit rates for subject D, and hit rates of valid-cued trials and correct rejection rates of invalid-cued trials for subject G.

The reaction time (RT) of valid-cued trials is defined as the time from the luminance change to the eye-movement initiation. The RT of the invalid-cued trials was similarly defined, although monkey G was trained to ignore the luminance change of the invalid cue, and the invalid trials in which monkey G made a response were false alarm trials. We only included trials in which the eye movement to the target were between 100 and 400 ms after the luminance change.

### Neurons’ selectivities

To determine how the value information was encoded by the neurons, linear regressions (*fitlm* function in Matlab Statistical Toolbox) were performed for each task variable for each individual unit in three task epochs:

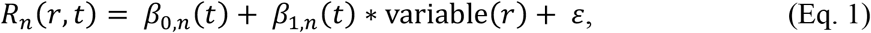

where *R_n_*(*r, t*) was the average neural response of unit *n* for a given trial *r* within time window *t*. In the single stimulus trials, the only relevant task variable is the stimulus value (*V_sin_*). In the double-stimuli trials, the task variables include the left stimulus value (*LV*), the right stimulus value (*RV*), the difference between the LV and the RV (*LV-RV*), the cued value (*CV*), the un-cued value (*UCV*), the difference between the CV and the UCV (*CV-UCV*), the salient value (*SV*), the non-salient value (*NSV*), the difference between the SV and the NSV (*SV-NSV*), the total value (*TV*) and the attention cue’s location (*att cue loc*). ε was independent Gaussian noise. The task epochs were the cue epoch (0.0 to 0.2 s after the cue onset), the cue-stimulus epoch (0.0 to 0.2 s after the stimulus onset), the early stimulus epoch (0.2 to 0.6 s after the stimulus onset), and the late stimulus epoch (0.0 to 0.4 s before the luminance change). A neuron is selective to a variable if *β_1,n_* is significantly different than 0 (p < 0.005).

A neuron is considered as value selective if it is selective to any value variables above during any task epochs (cue-stimulus, early stimulus and late stimulus). We further divide value selective neurons into positively tuned and negatively tuned neurons. Neurons with contradictory tuning are not included in **Figure 3, 5** and **Supplementary Table 1** (e.g., positively tuned to one value variable but negatively tuned to another value variable or positively tuned during one epoch but negatively tuned during another epoch).

### Decoding spatial attention

Linear discriminant analyses (LDA, *classify* function in Matlab Statistical Toolbox) were used to decode spatial attention location from the neuronal responses of LPFC neurons. LDA aims to classify an observation into one of the K classes through modeling the posterior probability p(Y = y|X = x), where X is the observation (neural responses), and Y is the predictor (left attention cue or right attention cue). The posterior probability is calculated through Bayes’ theorem which requires prior probability p(Y) and conditional probability density functions p(X = x|Y = y). LDA assumes that probability density function given each condition is normal distributed. Because left attention cue and right attention cue were equally likely, the prior was set to be 0.5.

Before conducting the LDA, we z-scored the responses of each neuron across all the trials and all time points (−400 to 600 ms around stimulus onset and −400 to 400 ms around luminance change). The normalized responses were further smoothed in 100 ms windows with a Gaussian kernel (sigma = 50 ms). The LDA was performed using a sliding window of 25 ms with 10 ms steps.

We constructed pseudo-neuron ensembles as follows. To balance the contribution from each neuron, we first selected the neurons with more than 100 trials in both the left attention cue condition and the right attention cue condition. We randomly chose 100 trials without replacements from each condition for each neuron and constructed the confusion matrix *X* ∈ ℝ^*M×T×N*^(with the resampled trials, where *M* is the number of trials (*M*=200, 100 trials in each cue condition), *T* is the number of time bins, and *N* is the number of neurons (subject D: *N*=102; subject G: *N*=93). To reduce noise, we ran the principal component analysis (PCA) and kept the first *P* components that captured at least 70% of the variance for the LDA.

To estimate how spatial attention signals fluctuated with time (**Figure 2c,e**), PCAs were run on the neuron dimension of *X* at each time bin. Only the first *P* principal components that captured at least 70% of the total variance were used to reconstruct the subspace (*Y* ∈ ℝ^*M×T×P*^). The decoder was trained and tested with the reconstructed subspace at each time bin. The posterior probability of attention cue location given the neuronal responses was used to quantify the decoder performance. The results were based on 200 independent leave-one-out cross validations. Significance was established by two-sided paired t-tests comparing the actual data against the posterior probabilities calculated in the same procedure but with the shuffled data.

In **Figure 2b, d**, a PCA was run on the neuron dimension of confusion matrix that combined all time bins (*X*′ ∈ ℝ^*MT×N*^). This ensured that the neural representation at each time bin shared the same subspace after PCA. Only the first *Q* principal components that captured at least 70% of the total variance were used to reconstruct the subspace (*Y*′ ∈ ℝ^*MT×Q*^). *Y*′ was reshaped into a ℝ^*M×T×Q*^ matrix to enable cross-temporal decoding. The decoder was trained with the neural responses in one time bin (**Figure 2b**: 25-ms time bin, stepped by 10 ms; **Figure 2d**: 50 to 200 ms before stimulus onset), and tested with the responses in all time bins.

### Linear Regression Models

In **Figure 4a, d**, linear regression models were fit to average firing rates calculated with a 25-ms time window, aligned to the stimulus onset and the luminance change, respectively. We constructed two linear regression models:

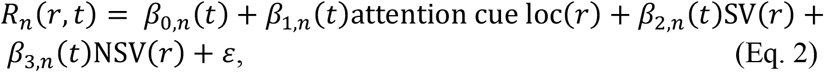

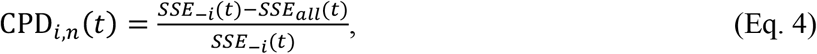

where *R_n_*(*r, t*) is the average firing rate of neuron n in trial r at time t, and the predictors were attention cue location (coded as −1 or 1), salient value (0, 1, 2, 4, or 8), non-salient value (0, 1, 2, 4, or 8), cued value (0, 1, 2, 4, or 8), un-cued value (0, 1, 2, 4, or 8). Each predictor was z-scored. Coefficient of partial determination was calculated as:

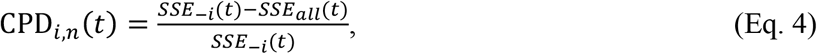

where CPD_*i,n*_(*t*) is the coefficient of partial determination of variable *i* in neuron *n*, *SSE_–i_*(*t*) is the residual sum of squares of the regression model without variable *i*, *SSE_all_*(*t*) is the residual sum of squares of the full model. The significance of CPD_*i,n*_(*t*) is tested against the baseline (paired-t test, p < 0.005 with FDR correction for multiple comparisons), which is the average CPD in a 200 ms time window before the attention cue onset:

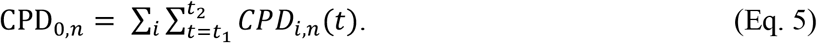

CPDs in **Figures 4b, d, e, f** were calculated similarly but with the average firing rate between the stimulus onset and the luminance change.

## Individual monkey analyses

All results presented in the main text are based on the combined data from both monkeys. Analyses based on each monkey individually can be found in the Supplementary Figures 2-6, corresponding to the Figures 1-5 in the main text. The results are consistent.

## Notes

### Competing Interest Statement

The authors have declared no competing interest.

