## Supplementary Figures & Table for "Bottom-up but not Top-down Attention Dominates the Value Representation in the Orbitofrontal Cortex"

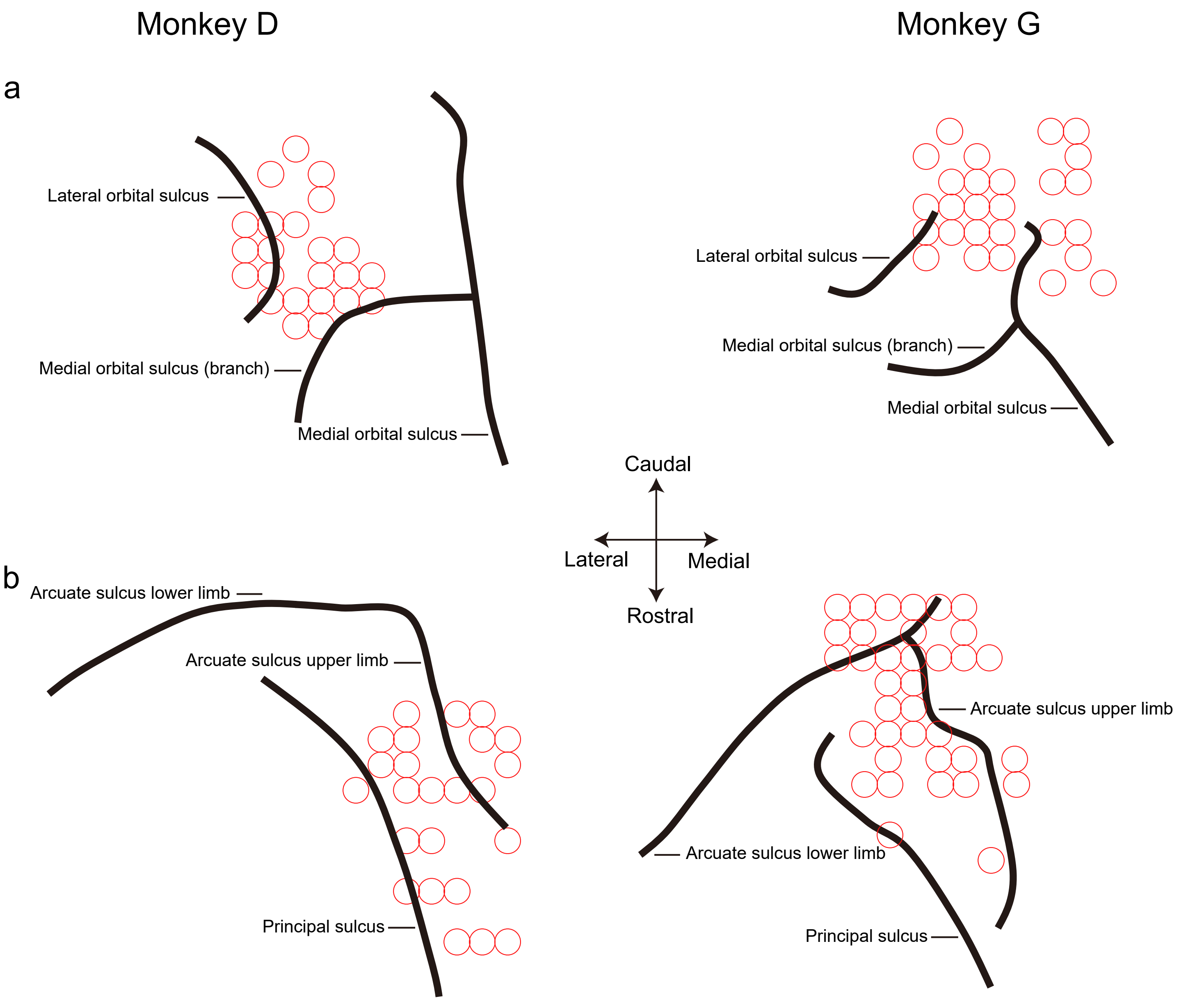


**Supplementary Figure 1**: The recording sites in **a.** the OFC and **b.** the DLPFC. Left column: monkey D; right column: monkey G. The circles indicate penetration location. The actual recording sites can be on the different side of a sulcus to what is illustrated here, as the penetrations can be angled.

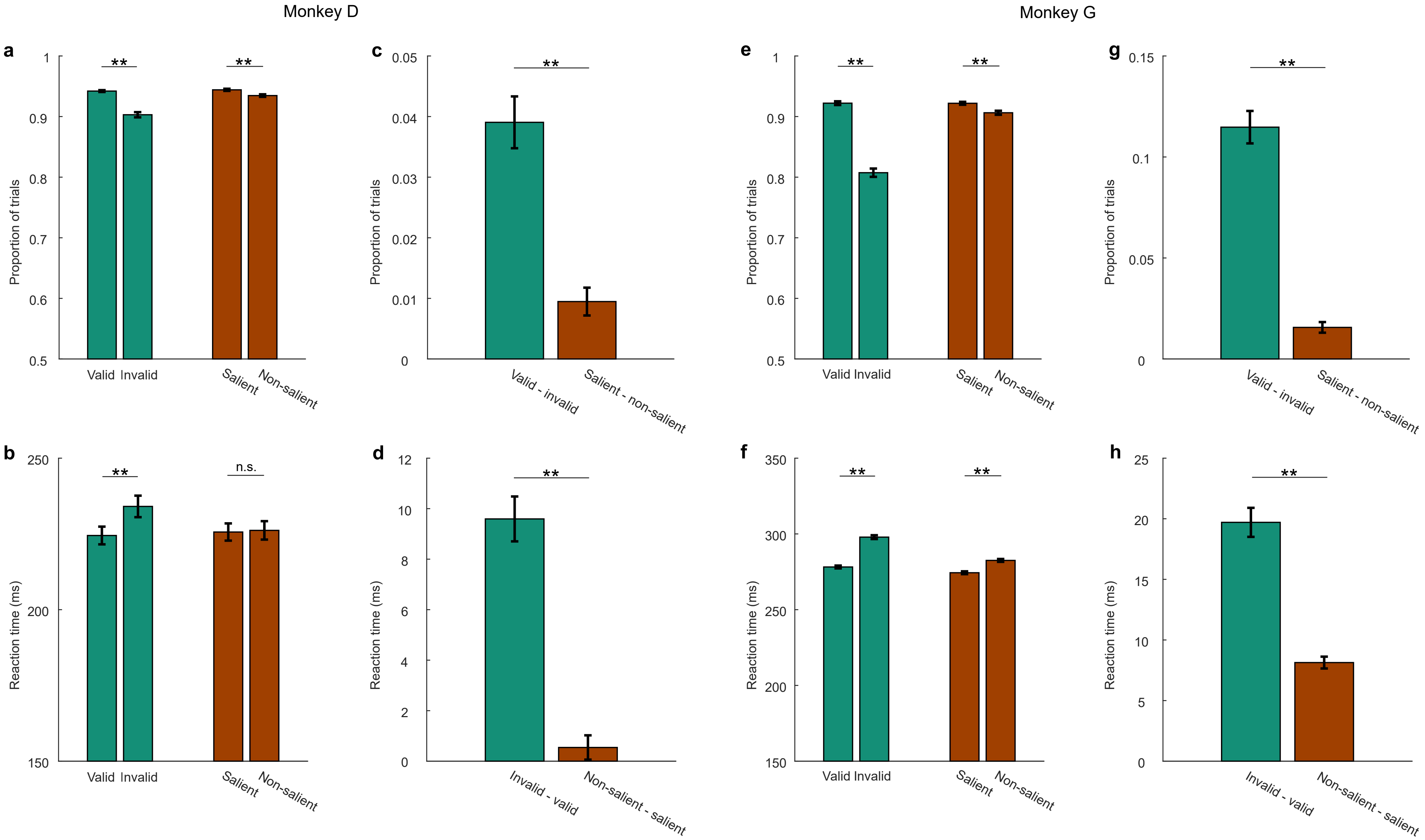


**Supplementary Figure 2**: The task performance of individual monkeys. Same convention as in **Figure 1**.

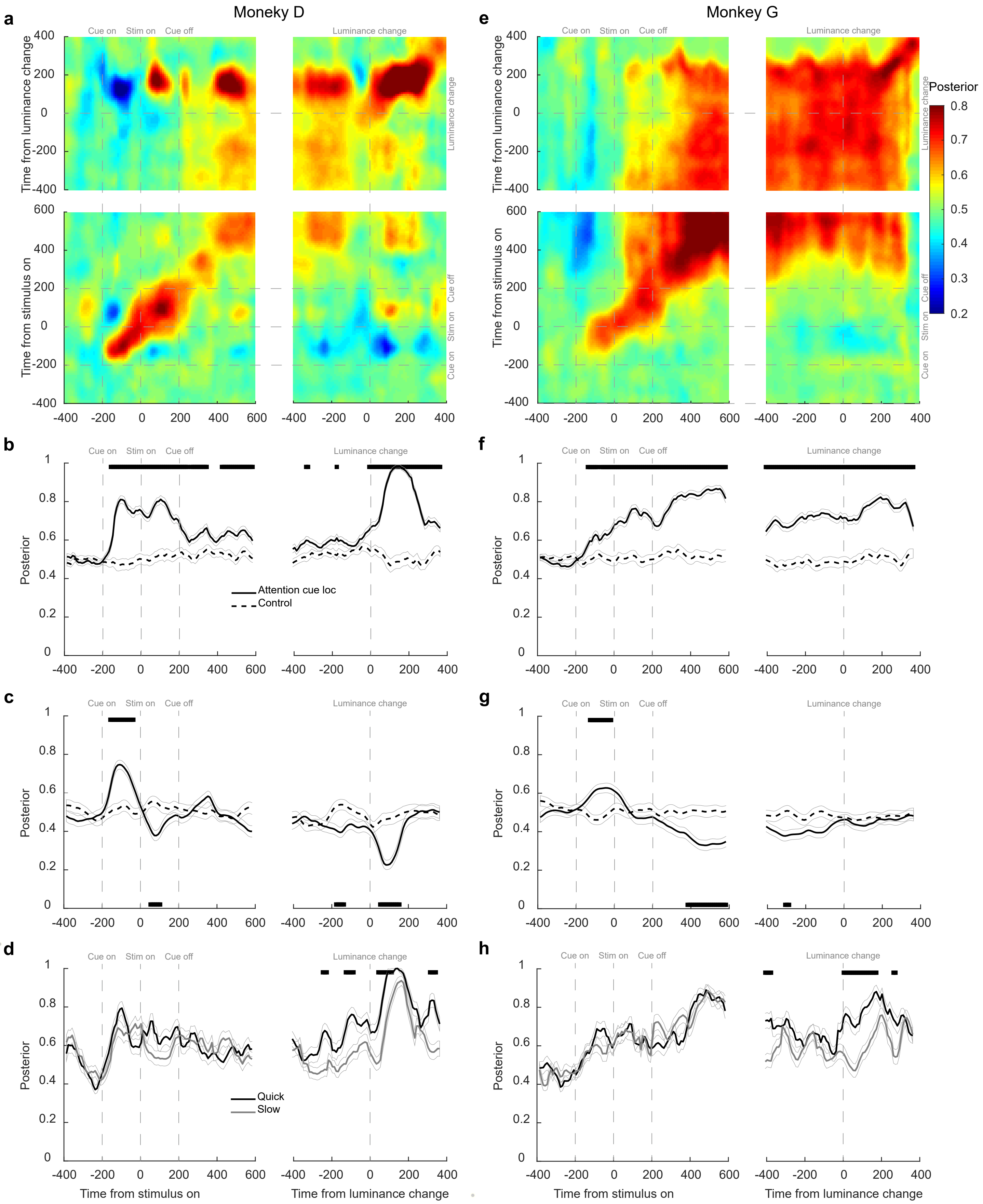


**Supplementary Figure 3**: LPFC encodes spatial attention in both monkeys. Same convention as **Figure 2**.

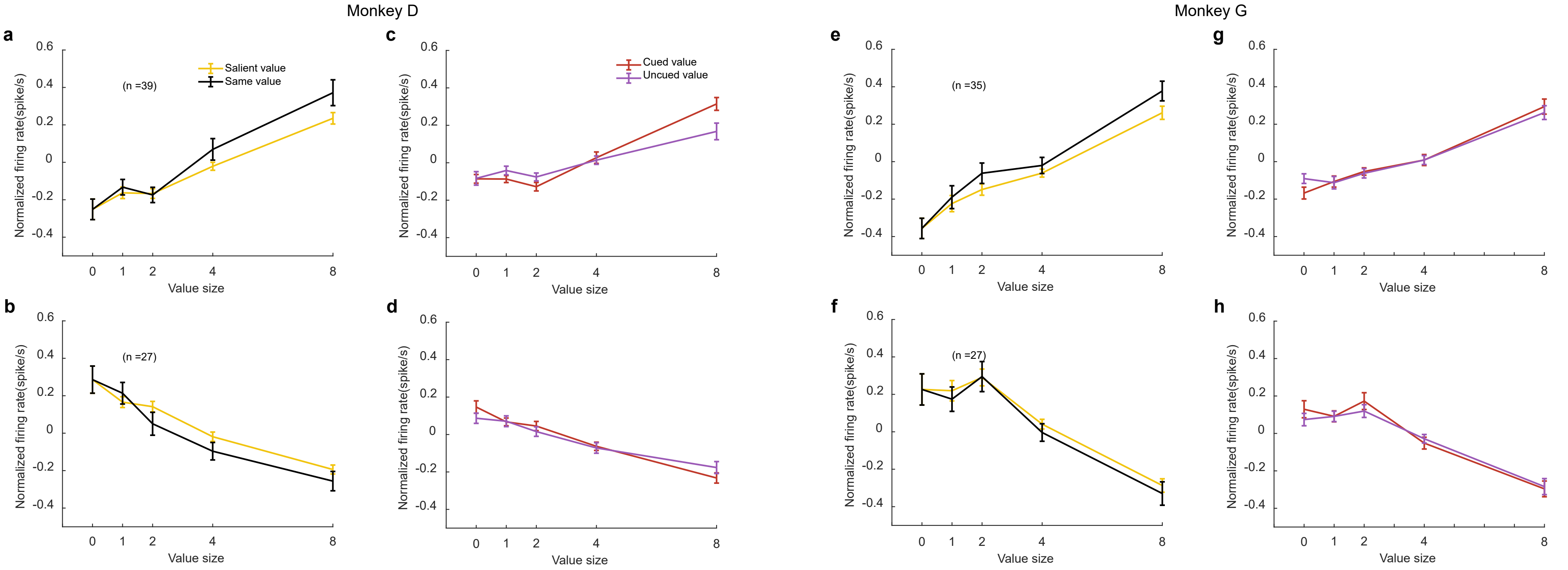


**Supplementary Figure 4**: Attentional modulation of the OFC responses in both monkeys. Same convention as in **Figure 3**.

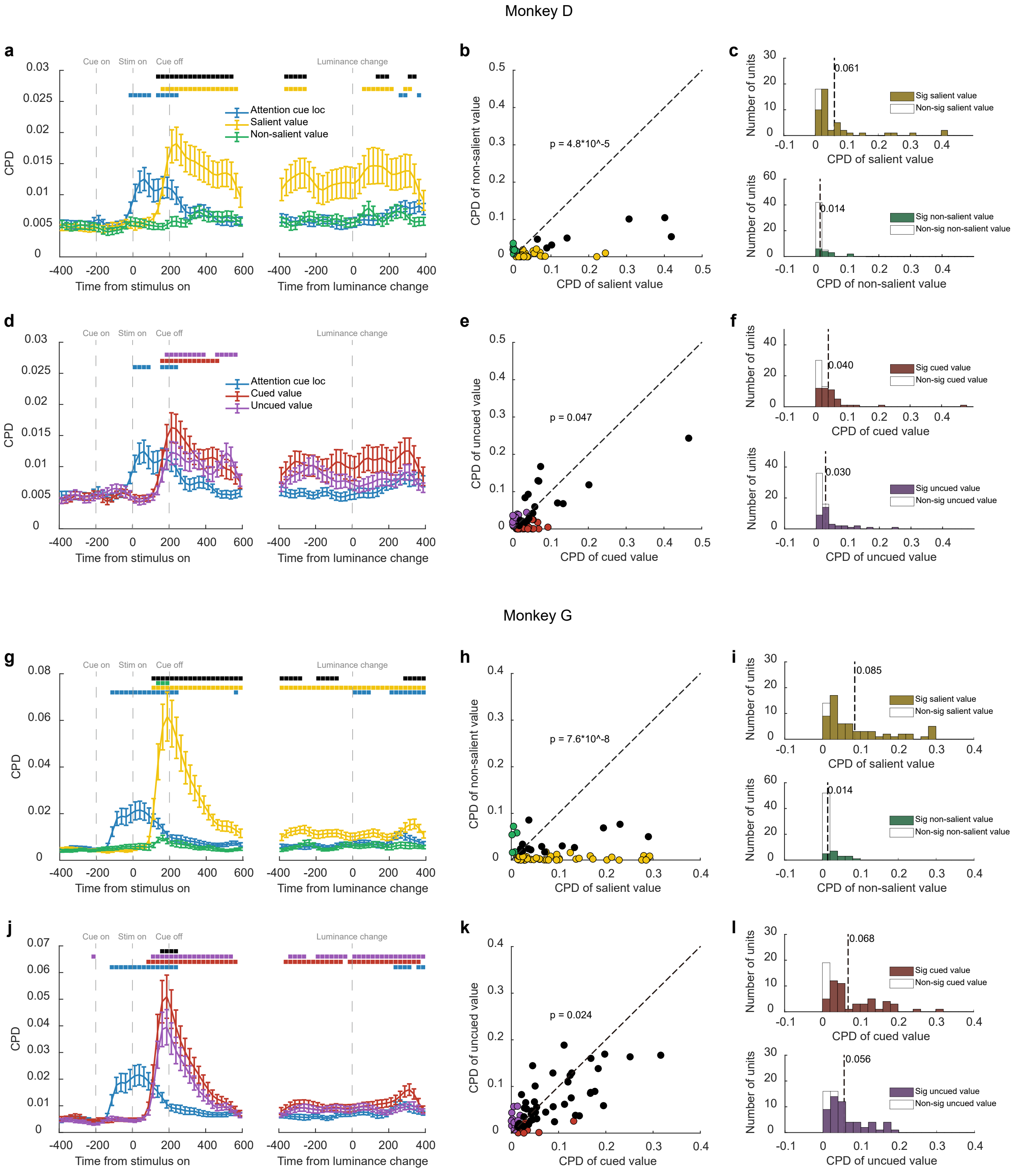


**Supplementary Figure 5**: OFC neuronal responses dominantly encoded the salient value in both monkeys. Same convention as in **Figure 4**.

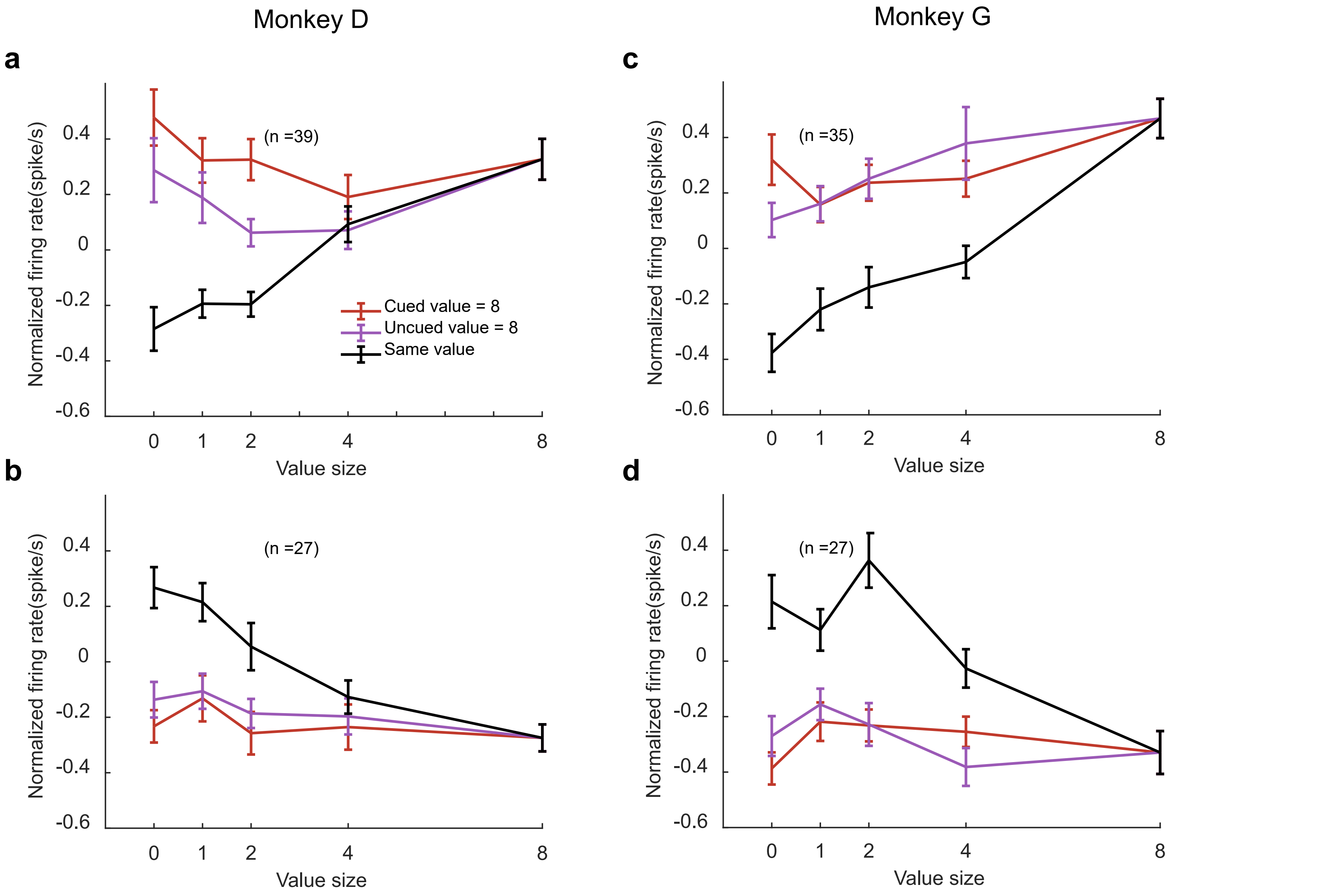


**Supplementary Figure 6**: Top-down attention failed to switch the value encoding in the OFC to the non-salient stimulus. in both monkeys. Same convention as in **Figure 5**.

**a** Both monkeys

Value selective OFC neurons (128/357)

| | $V_{sin}$ | LV | RV | LV-RV | CV | UCV | CV-UCV | SV | NSV | SV-NSV | TV | At least 1 var |
|---|---|---|---|---|---|---|---|---|---|---|---|---|
| Cue-stim | 5 | 9 | 18 | 1 | 14 | 8 | 2 | 20 | 10 | 10 | 19 | 35 |
| Early stim | 13 | 41 | 32 | 5 | 40 | 36 | 6 | 62 | 22 | 43 | 55 | 91 |
| Late stim | 4 | 26 | 16 | 2 | 21 | 24 | 6 | 40 | 16 | 18 | 44 | 69 |
| At least 1 epoch | 16 | 57 | 51 | 8 | 53 | 50 | 10 | 81 | 33 | 57 | 78 | 128 |

**b** Both monkeys

Correlation matrix.

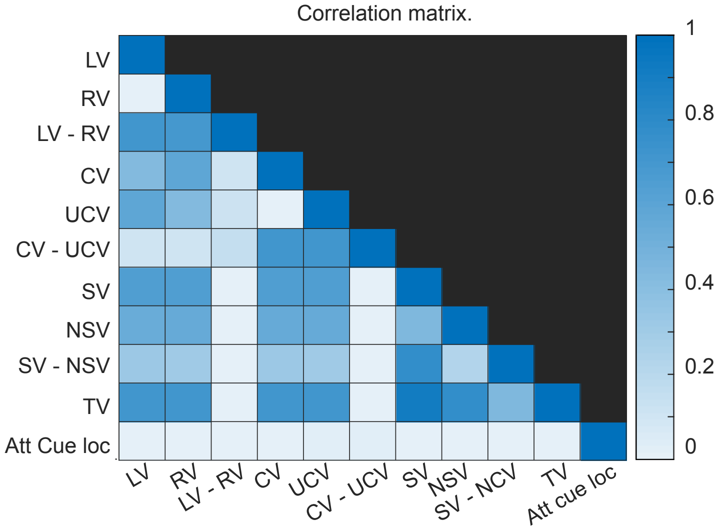


**c** Monkey D

Value selective OFC neurons (66/220)

| | $V_{sin}$ | LV | RV | LV-RV | CV | UCV | CV-UCV | SV | NSV | SV-NSV | TV | At least 1 var |
|---|---|---|---|---|---|---|---|---|---|---|---|---|
| Cue-stim | 1 | 4 | 2 | 1 | 2 | 3 | 1 | 3 | 1 | 2 | 3 | 11 |
| Early stim | 5 | 27 | 11 | 1 | 21 | 18 | 6 | 31 | 10 | 20 | 30 | 49 |
| Late stim | 2 | 15 | 7 | 1 | 12 | 12 | 5 | 22 | 10 | 9 | 24 | 39 |
| At least 1 epoch | 6 | 34 | 15 | 3 | 26 | 23 | 8 | 39 | 16 | 26 | 39 | 66 |

**d** Monkey G

Value selective OFC neurons (62/137)

| $V_{sin}$ | LV | RV | LV-RV | CV | UCV | CV-UCV | SV | NSV | SV-NSV | TV | At least 1 var |
|---|---|---|---|---|---|---|---|---|---|---|---|
| 4 | 5 | 16 | 0 | 12 | 5 | 1 | 17 | 9 | 8 | 16 | 24 |
| 8 | 14 | 21 | 4 | 19 | 18 | 0 | 31 | 12 | 23 | 25 | 42 |
| 2 | 11 | 9 | 1 | 9 | 12 | 1 | 18 | 6 | 9 | 20 | 30 |
| 10 | 23 | 36 | 5 | 27 | 27 | 2 | 42 | 17 | 31 | 39 | 62 |

**Supplementary Table 1**: Value-selective OFC neurons. **a**. Numbers of neurons selective to each value-related task variable (columns) during each epoch (rows). (linear regression, $H_0$: slope=0, $p < 0.005$). In the last row are the numbers of the neurons that showed selectivity to a given variable during at least one of the time windows. The definition of epochs is as follows. Cue-stim: 0.0 to 0.2 s after the stimulus onset (both the cue and the stimulus are on the screen); early stim: 0.2 to 0.6 s after the stimulus onset (only the stimulus on the screen); late stim: 0.0 to 0.4 s before the luminance change. In the last column are the number of neurons that show selectivity to at least one variable during a given time window. The task variables are: the stimulus value in single-stimulus trials ($V_{sin}$), the left stimulus value (*LV*), the right stimulus value (*RV*), the difference between the *LV* and the *RV* (*LV-RV*), the cued value (*CV*), the un-cued value (*UCV*), the difference between the *CV* and the *UCV* (*CV-UCV*), the salient value (*SV*), the non-salient value (*NSV*), the difference between the SV and the NSV (*SV-NSV*), and the total value (*TV*). **b.** Pearson correlation between different value variables. Calculated from all trials from the recording sessions. **c.** same as **a.**, but for Monkey D. **d.** same as **a.**, but for Monkey G.
